# Lipopolysaccharides derived from *Porphyromonas gingivalis* and *Escherichia coli:* differential and interactive effects on novelty-induced hyperlocomotion, blood cytokine levels and TRL4-related processes

**DOI:** 10.1101/2023.10.02.560531

**Authors:** Koji Saito, Yuri Aono, Arata Watanabe, Tetsuro Kono, Tomomi Hashizume-Takizawa, Hiroyuki Okada, Yasuhiro Kosuge, Hidenobu Senpuku, John L. Waddington, Tadashi Saigusa

**Affiliations:** Nihon University Graduate School of Dentistry at Matsudo, Oral Molecular Pharmacology, 2-870-1 Sakaecho-Nishi, Matsudo, Chiba 271-8587, Japan; Department of Pharmacology, Nihon University School of Dentistry at Matsudo, 2-870-1 Sakaecho-Nishi, Matsudo, Chiba 271-8587, Japan; Department of Histology, Nihon University School of Dentistry at Matsudo, 2-870-1 Sakaecho-Nishi, Matsudo, Chiba 271-8587, Japan; Department of Microbiology and Immunology, Nihon University School of Dentistry at Matsudo, 2-870-1 Sakaecho-Nishi, Matsudo, Chiba 271-8587, Japan; Laboratory of Pharmacology, Nihon University School of Pharmacy, 7-7-1 Narashinodai, Funabashi, Chiba 274-8555, Japan; School of Pharmacy and Biomolecular Sciences, RCSI University of Medicine and Health Sciences, St. Stephen’s Green, Dublin 2, Ireland

**Keywords:** *Porphyromonas gingivalis*, lipopolysaccharide, locomotion, Toll-like receptor, IL-10, mouse

## Abstract

Lipopolysaccharide (LPS), a component of the Gram-negative bacterial cell wall, activates Toll-like receptors (TLRs). *Porphyromonas gingivalis* (*Pg*) may be involved in the progression of periodontal disease. Mice exposed to a novel environment show hyperlocomotion that is inhibited by systemic administration of LPS derived from *Escherichia coli* (*Ec*-LPS). However, whether *Pg*-LPS influences novelty-induced locomotion is unknown. Accordingly, we carried out an open field test to analyse the effects of *Pg*-LPS. For comparison, effects of *Ec*-LPS were also studied. We also investigated the influence of systemic administration of *Pg*-LPS or *Ec*-LPS on IL-6, TNF-alpha, and IL-10 levels in blood, as they could be involved in the changes in locomotion. The TLR4 receptor antagonist TAK-242 was used to study the involvement of TLR4. Since *Pg*-LPS may block TLR4 *in vitro*, we analysed the effects of *Pg*-LPS on *Ec*-LPS-induced changes in behavioural and biochemical parameters. Male ddY mice were used. Compounds were administered intraperitoneally. *Ec*-LPS (840 µg/kg), but not *Pg*-LPS (100, 500 and 840 µg/kg), inhibited novelty-induced locomotion, which was reversed by TAK-242 (3.0 mg/kg). *Ec*-LPS (840 µg/kg) increased blood levels of IL-6 and IL-10, which was antagonized by TAK-242 (3.0 mg/kg). However, TAK-242 did not inhibit *Ec*-LPS-induced increases in TNF-alpha levels in blood. *Pg*-LPS (100, 500, and 840 µg/kg) did not alter blood IL-6, TNF-alpha, or IL-10 levels. The *Ec*-LPS-induced increase in blood IL-10, but not IL-6 and TNF-alpha, levels was inhibited by *Pg*-LPS (500 µg/kg). These results suggest that TLR4 stimulation mediates the inhibition of novel environment-induced locomotion in mice following systemic administration of *Ec*-LPS, while also increasing blood IL-6 and IL-10 levels. In contrast, *Pg*-LPS did not exhibit these effects. The present study also provides *in vivo* evidence that *Pg*-LPS can inhibit TLR4-mediated increases in blood IL-10 levels, which is thought to prevent the development of periodontal disease.

## Introduction

*Porphyromonas gingivalis* (*Pg*) is a Gram-negative, rod-shaped bacterium that may be involved in the progression of periodontal disease. Lipopolysaccharide (LPS), a component of the Gram-negative bacterial cell wall derived from *Pg* (*Pg*-LPS), has been implicated as a key molecule in the development of periodontitis.

*In vitro* studies using cultured cells from periodontal tissue have revealed *Pg*-LPS to induce effects distinct from LPS derived from the prototype Gram-negative, rod-shaped bacterium *Escherichia coli* (*Ec*-LPS). Notably, increases in interleukin (IL)-6 expression observed in gingival fibroblasts [1] and periodontal ligament cells [2] following treatment with *Pg*-LPS were reduced relative to those following treatment with *Ec*-LPS. Additionally, treatment with *Pg*-LPS failed to alter IL-6 and tumor necrosis factor (TNF)-alpha levels significantly in dental pulp stem cells, while *Ec*-LPS treatment markedly increased these cytokines [3]. Moreover, treatments with *Pg*-LPS had no significant effect on cytokine production in monocyte-derived dendritic cells, whereas *Ec*-LPS treatment led to a significant increase in IL-6 and IL-10 production [4].

Despite these observed differences in cytokine responses, the specific mechanisms underlying the divergent effects of *Pg*-LPS and *Ec*-LPS on cytokine production remain unclear. Nevertheless, it is plausible that variations in receptors interacting with *Pg*-LPS and *Ec*-LPS may contribute to these discrepancies. Toll-like receptors (TLRs) are well-known pattern recognition sites that bind pathogen-associated molecules, ultimately activating immunity and host defense [5]. Notably, LPS stimulates a subtype of TLR known as TLR4, which is expressed on the plasma membrane and mediates various processes, including cytokine synthesis and secretion. Studies using cultured monocytic cells have suggested that increases in IL-6 and TNF-alpha levels induced by *Ec*-LPS are mediated via TLR4 activation. On the other hand, the effects of *Pg*-LPS on IL-6 and TNF-alpha were proposed to be mediated via TLR4 and/or another subtype of TLR known as TLR2 [6]. Moreover, differences in the effects of *Pg*-LPS and *Ec*-LPS can be attributed to their differential abilities to activate TLR4; previous research has shown that *Ec*-LPS strongly stimulates TLR4, whereas *Pg*-LPS weakly stimulates [7–10] or blocks [11–14] TLR4. *In vivo* studies in rats have demonstrated that *Pg*-LPS induces effects on extracellular cytokine levels that differ from those induced by *Ec*-LPS. For example, our studies using urethane-anesthetized rats indicated that intra-gingival administration of *Ec*-LPS failed to alter cytokine levels, while *Pg*-LPS increased gingival extracellular levels of TNF-alpha [15].

Systemic administration of *Ec*-LPS in mice is known to induce a variety of sickness behaviors, such as reductions in water and food intake and loss of body weight [16]. These behavioral changes are suggested to be mediated by increases in the synthesis and release of cytokines that can be detected in the systemic circulation [16]. Notably, the sickness behavior induced by systemic administration of *Ec*-LPS also includes reduction in locomotor activity [17]. However, it remains unknown whether systemic administration of *Pg*-LPS in mice produces inhibitory effects on locomotion similar to those induced by *Ec*-LPS. Therefore, we conducted an open field test to examine whether a single systemic administration of *Pg*-LPS in mice affects locomotor activity in a novel environment, to further characterize the effects of *Pg*-LPS *in vivo*. For comparison, we also analyzed the effects of *Ec*-LPS on novelty-induced increases in locomotor activity under the same experimental conditions. Additionally, we investigated the influence of systemic administration of *Pg*-LPS or *Ec*-LPS on blood levels of cytokines, particularly IL-6, TNF-alpha and IL-10, as these cytokines may have pro-inflammatory (IL-6 and TNF-alpha) or anti-inflammatory (IL-10) effects that could be involved in the observed changes in mouse behavior. Furthermore, animal experiments have shown that systemic inflammation induces increases in spleen mass [18]. For example, repeated oral administration of *Pg* has been found to increase spleen weight in mice [19]. The spleen plays a crucial role in modulating immune responses by promoting the differentiation and activation of T and B cells. Thus, we investigated the influence of systemic administration of *Pg*-LPS or *Ec*-LPS on spleen weight, together with the numbers of T cells and B cells using flow cytometry. To elucidate the involvement of TLR4 in the effects of LPS on behavioral and biochemical parameters in mice, we co-administered TAK-242, a selective TLR4 receptor antagonist [20]. Given the *in vitro* experiments suggesting that *Pg*-LPS may exhibit partial agonist- or antagonist-like properties on TLR4, as discussed above [7–14], we proceeded to analyse the effects of *Pg*-LPS on *Ec*-LPS-induced changes in behavioural and biochemical parameters in mice.

Preliminary results from these investigations have been presented at the Annual Meeting of the Japanese Pharmacological Society, 2022, and the Annual Meeting of the Japanese Association for Oral Biology, 2023.

## Materials and Methods

### Animals

Male ddY mice (Sankyo Laboratory Service Co. Ltd., Tokyo, Japan), weighing between 25 and 30 g were used. These were kept at constant room temperature (23 ± 2 °C) and relative humidity (55 ± 5%) under a 12 h day:night cycle (light on: 07.00 a.m.), with ad libitum access to food and water. All experiments were approved by the Animal Experimentation Committee of Nihon University School of Dentistry at Matsudo (AP19MAS015) and were performed in accordance with national and international guidelines for the care and welfare of animals. All efforts were made to minimise animal suffering and to reduce the number of animals used.

### Drugs and treatments

Commercially available LPS derived from *Ec* and *Pg*, namely *Ec*-LPS (O55:B5; Sigma-Aldrich, St. Louis, MO, USA) and *Pg*-LPS (LPS-PG Ultrapure; InvivoGen, San Diego, CA, USA) were used. They were diluted with saline and aliquots were stored at -20℃ prior to use within two months. TAK-242 (Sigma-Aldrich) was used as a TLR4 antagonist. It was diluted with saline and a small amount of dimethyl sulfoxide (<0.1%) was added immediately prior to use. The doses of these compounds were determined by a series of pilot experiments based on previous behavioural studies (LPS: [17, 21]; TAK-242: [20]).

Compounds were injected i.p. in a volume of 0.1 ml per 10 g body weight, with each mouse used only once. Mice were treated with LPS or its corresponding vehicle 4 h prior to the open field test and the subsequent measurements of spleen weight, flow cytometry, and blood cytokine levels. TAK-242 or its vehicle was administered 1 h before designated procedures.

### Open field test

On the day of experiments, each mouse was individually placed in a plastic cage (length × width × height: 32 × 21 × 13 cm), the floor of which was covered with sawdust to a depth of approximately 2 cm, and habituated to these conditions for at least 60 min. Then, mice treated with LPS and/or TAK-242 or their corresponding vehicles (see Drugs and treatments) was transferred to a Plexiglas box (length × width × height: 30 × 30 × 20 cm) that was slightly larger and lacked floor sawdust. Locomotion in the novel environment was recorded for a period of 30 min via a video camera located above the Plexiglas box and total distance traveled (cm) was determined by an automated system for behavioural assessment (SMART, Panlab, Barcelona, Spain). In designated experiments we carried out additional analyses of locomotion in the center (20 × 20 cm) *vs* the periphery (within 5 cm of each wall) of the Plexiglas box, since avoiding the center zone of the novel environment is regarded as a sign of anxiety that could be elicited by systemic administration of LPS. After evaluation of each mouse the Plexiglas box was cleaned with ethanol followed by water before proceeding to the next mouse.

### Measurement of spleen weight and flow cytometry

After treatments with LPS and/or TAK-242 or their corresponding vehicles (see Drugs and treatments) mice were euthanized by isoflurane (5%) inhalation. Their spleens were collected and weighed by a precision electronic weighing scale and single-cell suspensions were prepared in saline using a homogenizer.

Surface markers of splenocytes were identified using monoclonal antibodies in conjunction with two immunofluorescence analysis with a flow cytometer (BD Accuri C6 Plus: BD, Mountain View, CA, USA). Antibodies used were Fluoroscein isothiocyanate (FITC)-conjugated anti-mouse CD4 and CD21/CD35 (CR2/CR1: Biolegend, San Diego, CA, USA) and Phycoerythrin (PE)-conjugated anti-mouse CD3 (17A2: Proteintech, Rosemont, IL, USA). Isotype controls used were FITC-labelled Rat IgG2b Isotype Control (LTF-2) (Proteintech, Rosemont, IL, USA) and PE-labelled Rat IgG2b Isotype Control (Becton Dickinson Pharmingen, San Jose, CA, USA). Double-labelled surface phenotypes were CD4/CD3 and CD3/CD21. Cells were pre-incubated with anti-CD16/CD32 antibody to block nonspecific antibody binding. Then, cells were incubated with the above specific antibodies for 30 min and washed twice in 1% bovine serum albumin in phosphate buffered saline. Seven-amino-actinomycin D (Biolegend, San Diego, CA, USA) was added before analysis with a flow cytometer to exclude nonviable cells. Data were analyzed using commercial software (FlowJo, BD Biosciences, Franklin Lakes, NJ, USA).

### Measurement of blood cytokine levels

A disposable lancet (4 mm, Goldenrold Animal Lancet, MEDIpoint, Mineola NY, USA) was used to take blood samples from the submandibular vein of mice treated with LPS and/or TAK-242 or their corresponding vehicle (see Drugs and treatments). Approximately 0.5 ml of blood was quickly collected without total anesthesia [22]. Blood was allowed to coagulate for 10-15 min at room temperature, then centrifuged at 1900 × g for 10 min at 4℃, with separated sera then stored at - 70℃ until further analysis.

Levels of IL-6, TNF-alpha and IL-10 in serum were determined using bead-based Multi-Plex kits (MILLIPLEX^®^ Mouse Cytokine/Chemokine Magnetic Bead Panel 96-Well Plate Assay, Merck KGaA, Cat. No. MCYTOMAG-70K-03, Rockland, MA, USA). In order to remove particulates, samples were centrifuged at 16000 × g for 4 min at 4℃ before assay and diluted 2-fold using medium provided by the kit manufacture. Twenty-five µl of diluted samples were then transferred into a 96-well plate and incubated overnight at 4℃ with shaking to immobilize antibody beads. Liquid was removed, followed by two washes, and detection beads were then added into each well. The plate was then incubated for 1h at room temperature, followed by addition of streptavidin-phycoverthrin for 30 min with shaking. After the supernatant was removed, 150 µl of drive fluid to resuspend the beads was added to each well and the plate was read using MAGPIX plate reader with xPOINT software (Luminex® 100/200™ System, Luminex Corp., Austin, TX, USA). Data were analyzed using MILLIPLEX Analyst software.

### Statistical analysis

All values are expressed as mean and S.E.M. Comparisons of (1) effects of various doses of *Pg*- and *Ec*-LPS on distance travelled and (2) effects of TAK-242 or *Pg*-LPS on *Ec*-LPS-induced changes in distance travelled, body weight, spleen weight, cellular components of spleen and blood cytokine levels were performed using one-way ANOVA followed by post hoc Scheffé’s test where appropriate. Effects of *Ec*-LPS on distance travelled in central *vs* peripheral zones of the novel environment were compared using Student’s *t*-test. Statistical significance was considered to be *P* < 0.05.

## Results

### *Ec*-LPS but not *Pg*-LPS inhibits novelty-induced locomotor activity

Locomotion in mice treated with vehicle and exposed to the open field was not altered by *Pg*-LPS (100, 500 or 840 µg/kg). In contrast, such locomotion was inhibited by *Ec*-LPS (Fig. 1: one-way ANOVA, *F* (3, 25) = 4.11, *P* < 0.05). Post hoc Sheffé’s test revealed that the effects of 840 µg/kg *Ec*-LPS differed significantly from those of vehicle (*P* < 0.05).

**Fig 1.**
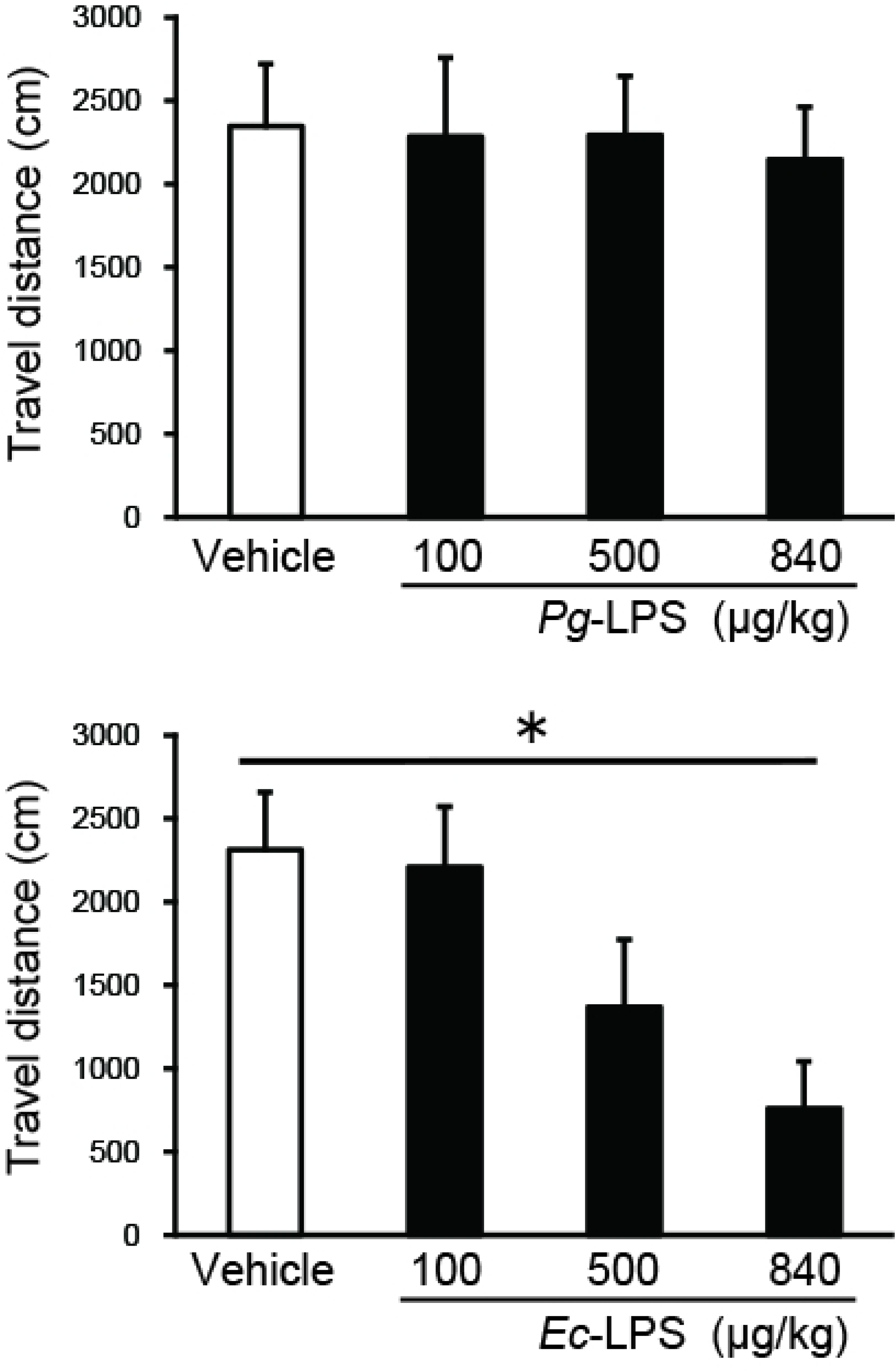
Upper panel: Effects of intra-peritoneal injection of *Pg*-LPS on novel environment-induced increases in locomotor activity in mice. Lower panel: Effects of intra-peritoneal injection of *Ec*-LPS on novel environment-induced increases in locomotor activity in mice. Vertical bars indicate S.E.M., n = 7-8 per group. \**P* < 0.05, *Pg*- or *Ec*-LPS *vs* vehicle.

### *Ec*-LPS-induced inhibition of novelty-induced locomotor activity is antagonised by TAK-242 but not by *Pg*-LPS

Pretreatment with the TLR4 antagonist TAK-242 (3.0 mg/kg), which did not alter locomotion when given alone, antagonized the inhibition of locomotion induced by 840 µg/kg *Ec*-LPS (Fig. 2: one-way ANOVA, *F* (3, 22) = 7.03, *P* < 0.01). Post hoc Sheffé’s test revealed that the effect of 840 µg/kg *Ec*-LPS pretreated with 3.0 mg/kg TAK-242 differed significantly from the effect of 840 µg/kg *Ec*-LPS pretreated with vehicle (*P* < 0.05). Pretreatment with *Pg*-LPS (500 µg/kg), which did not alter locomotion when given alone, did not significantly influence the inhibition of locomotion induced by 840 µg/kg *Ec*-LPS.

**Fig 2.**
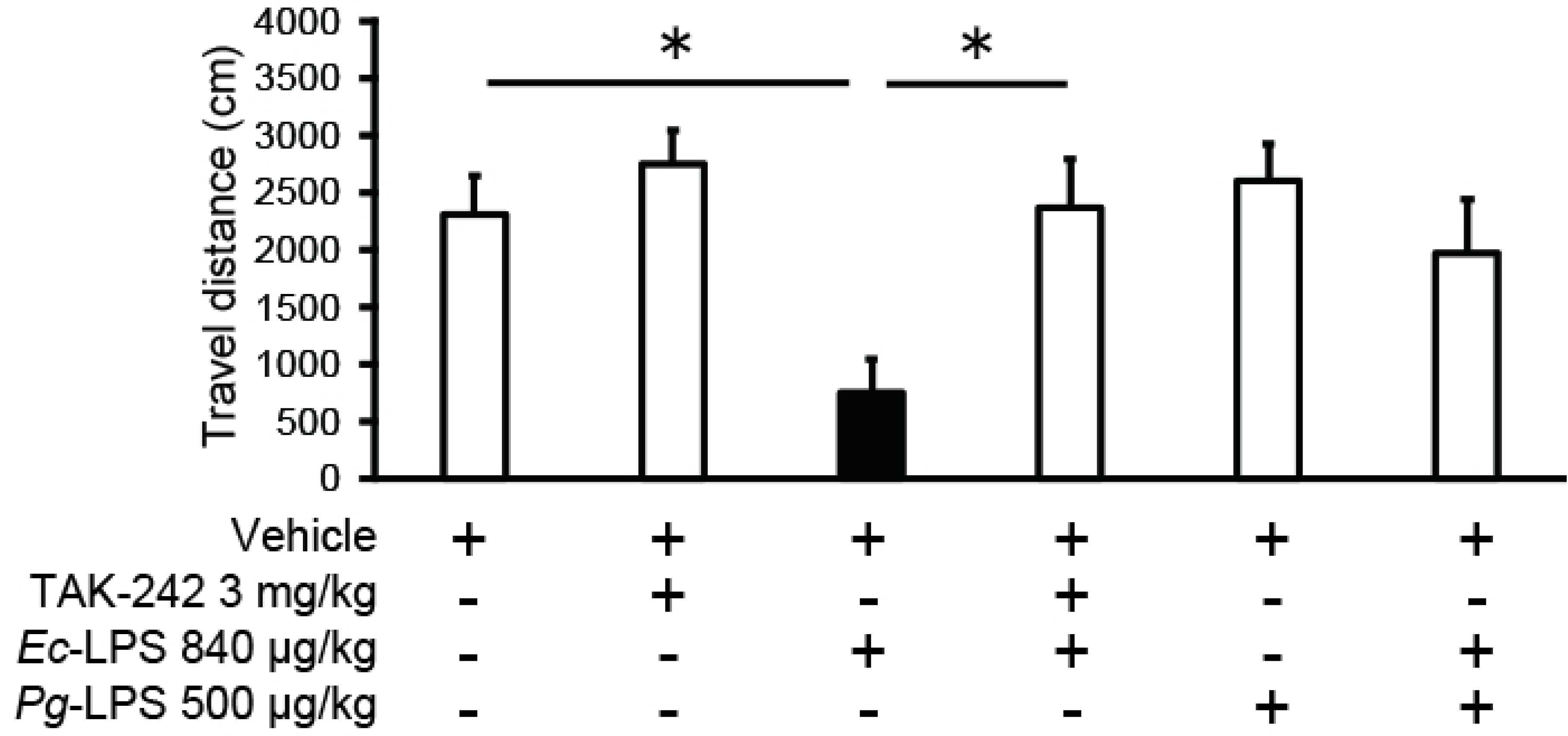
Effects of TAK-242 or *Pg*-LPS on *Ec*-LPS-induced reduction in novelty-induced locomotor activity in mice. Vertical bars indicate S.E.M., n = 6-7 per group. \**P* < 0.05 *vs Ec*-LPS + vehicle.

### *Ec*-LPS-induced inhibition of novelty-induced locomotion is evident in both the central and peripheral zones

As differential effects on locomotion in the peripheral *vs* central zones of the open field implicate anxiety-/depression-related processes [23], we analyzed the effects of *Ec*-LPS (840 µg/kg) on locomotion in these two zones. *Ec*-LPS treatment inhibited locomotion in both the central and peripheral zones (Fig. 3: Student’s *t*-tests, *Ec*-LPS 840 µg/kg *vs* vehicle: central zone: *t* (12) = 2.19, *P* < 0.05; peripheral zone: *t* (12) = 3.93, *P* < 0.01).

**Fig 3.**
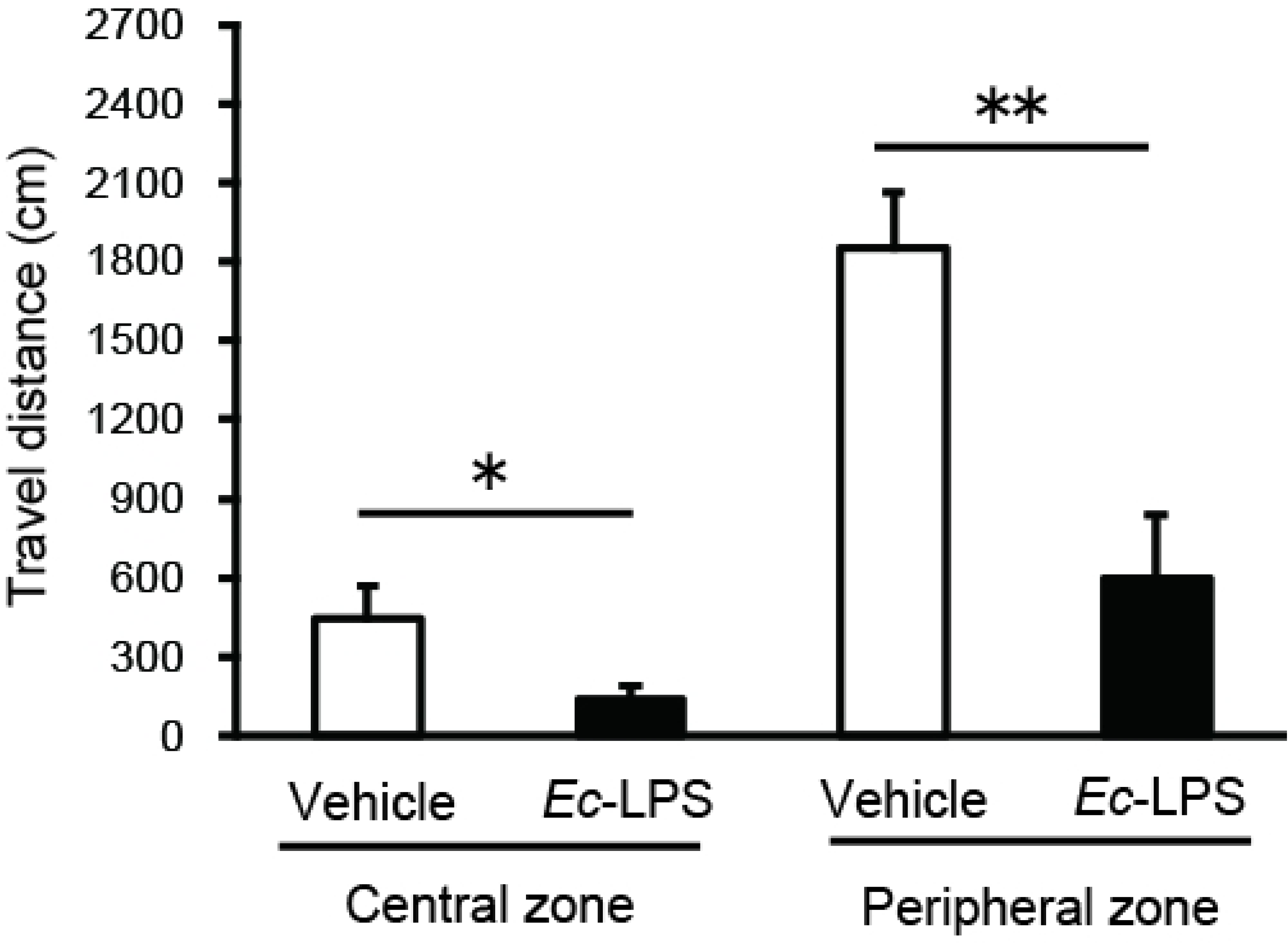
Effects of intra-peritoneal injection of *Ec*-LPS on novel environment-induced increases in locomotor activity within central and peripheral zones of the open field in mice. Vertical bars indicate S.E.M., n = 7 per group. \**P* < 0.05, ** *P* < 0.01, *Ec*-LPS *vs* vehicle.

### *Ec*- and *Pg*-LPS treatments, with or without TAK-242 pretreatment, fail to alter spleen weight and T or B cell subsets

We measured the body and spleen weight of mice without any treatment, vehicle alone, TAK-242 (3.0 mg/kg), *Pg*-LPS (500 or 840 μg/kg), *Ec*-LPS (840 μg/kg) alone, TAK242 (3.0 mg/kg) followed by administration of *Ec*-LPS (840 μg/kg), *Pg*-LPS (500 μg/kg) followed by administration of *Ec*-LPS (840 μg/kg). Though there appeared to be some numerical variation, neither body nor spleen weight differed significantly across these groups (Table 1).

**Table 1.** Body weight (g) and spleen weight (mg) of mice receiving LPS and/or TAK242 treatments.

|  | Body weight<br>(g) | Spleen<br>weight<br>(mg) |
| --- | --- | --- |
| Without treatment | 23.0 ± 0.0 | 125.8 ± 9.7 |
| Vehicle | 23.0 ± 0.0 | 136.7 ± 13.6 |
| TAK-242 (3.0 mg/kg) | 24.8 ± 1.3 | 130.8 ± 8.7 |
| *Pg*-LPS (500 µg/kg) | 22.3 ± 0.3 | 105.5 ± 2.4 |
| *Pg*-LPS (840 µg/kg) | 21.0 ± 0.0 | 147.0 ± 25.1 |
| *Ec*-LPS (840 µg/kg) | 22.0 ± 0.0 | 114.0 ± 3.0 |
| TAK-242 (3.0 mg/kg) + *Ec*-LPS (840 µg/kg) | 24.4 ± 1.2 | 132.6 ± 4.2 |
| *Pg*-LPS (500 µg/kg) + *Ec*-LPS (840 µg/kg) | 23.5 ± 2.1 | 145.4 ± 11.8 |

Mean and S.E.M., n = 3-6 per group.

Next, we assessed the cellular composition of the spleen in these same experimental groups by flow cytometry and determined the numbers of CD4^+^ helper T cell and CD21^+^ mature B cell subsets. Though there appeared to be some numerical variation, neither the number of CD4^+^ T cells nor the number of CD21^+^ B cells differed significantly across these groups (Table 2).

**Table 2.** Cellular components of spleen in mice receiving LPS and/or TAK242 treatments.

|  | CD4<sup>+</sup> T cell<br>(× 10<sup>8</sup>) | CD21<sup>+</sup> B cell<br>(× 10<sup>8</sup>) |
| --- | --- | --- |
| Without treatment | 5.3 ± 0.7 | 24.4 ± 3.5 |
| Vehicle | 5.7 ± 0.5 | 30.6 ± 7.1 |
| TAK-242 (3.0 mg/kg) | 7.2 ± 1.3 | 19.9 ± 5.2 |
| *Pg*-LPS (500 µg/kg) | 8.1 ± 1.0 | 10.9 ± 1.3 |
| *Pg*-LPS (840 µg/kg) | 5.1 ± 0.7 | 23.7 ± 5.5 |
| *Ec*-LPS (840 µg/kg) | 4.5 ± 0.6 | 22.0 ± 1.9 |
| TAK-242 (3.0 mg/kg) + *Ec*-LPS (840 µg/kg) | 9.7 ± 1.4 | 27.8 ± 2.7 |
| *Pg*-LPS (500 µg/kg) + *Ec*-LPS (840 µg/kg) | 10.8 ± 2.6 | 25.8 ± 2.9 |

Mean and S.E.M., n = 3-6 per group.

### *Ec*-LPS, but not *Pg*-LPS, increases blood IL-6 levels that are antagonized by TAK-242 but not by *Pg*-LPS

Baseline IL-6 levels in blood samples were 20.0 ± 11.1 pg/ml (n = 6) and these levels were not altered by administration of vehicle (Fig. 4). *Ec*-LPS (840 µg/kg), but not *Pg*-LPS (840 µg/kg), increased blood levels of IL-6 (Fig. 4: one-way ANOVA, *F* (7, 34) = 8.51, *P* < 0.001). This *Ec*-LPS-induced increase in blood level of IL-6 was antagonized by pretreatment with TAK-242 (3.0 mg/kg) that did not influence blood IL-6 levels when given alone (Fig. 4). *Pg*-LPS (500 µg/kg) pretreatment failed to alter either baseline IL-6 or *Ec*-LPS (840 µg/kg)-induced increases in IL-6. Post hoc Sheffé’s tests revealed that the effects of *Ec*-LPS (840 µg/kg) differed significantly from those of vehicle (*P* < 0.01), 500 µg/kg *Pg*-LPS (*P* < 0.01), 840 µg/kg *Pg*-LPS (*P* < 0.01) and 3.0 mg/kg TAK-242 + 840 µg/kg *Ec*-LPS (*P* < 0.05).

**Fig 4.**
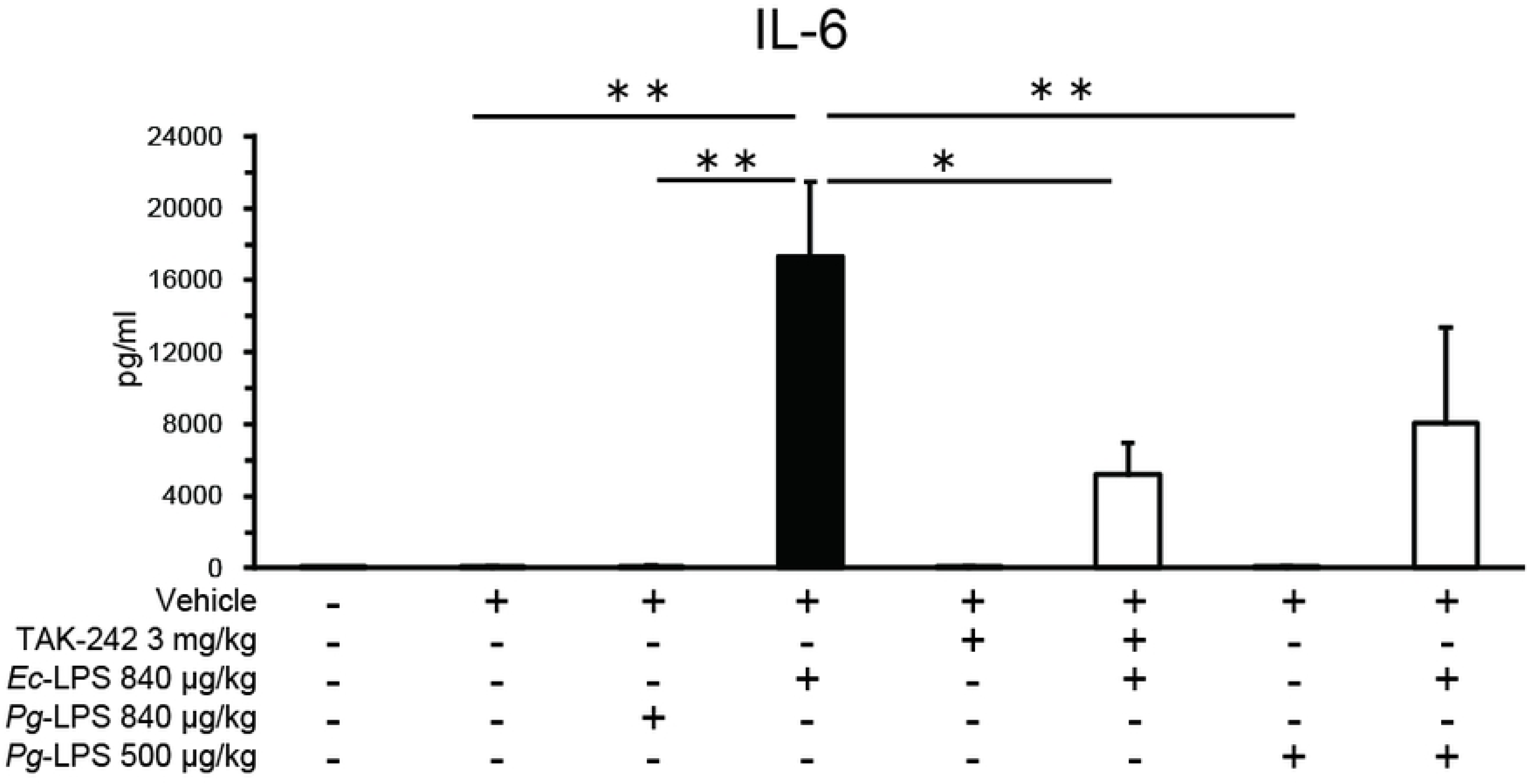
Effects of intra-peritoneal injection of *Pg*- or *Ec*-LPS with TAK-242 and of co-administration of *Pg*- and *Ec*-LPS on IL-6 levels (pg/ml) in blood samples of mice. Vertical bars indicate S.E.M., n = 4-6 per group. \**P* < 0.05, ** *P* < 0.01 *vs Ec*-LPS + vehicle.

### *Ec*-LPS, but not *Pg*-LPS, increases blood TNF-alpha levels that are not antagonized by TAK-242 or *Pg*-LPS

Baseline TNF-alpha levels in blood samples were 7.2 ± 1.2 pg/ml (n = 6) and these levels were not altered by administration of vehicle (Fig. 5). *Ec*-LPS (840 µg/kg), but not *Pg*-LPS (840 ug/kg), increased blood levels of TNF-alpha (Fig. 5: one-way ANOVA, *F* (7, 34) = 9.36, *P* < 0.001). This *Ec*-LPS-induced increase in blood levels of TNF-alpha was not influenced by either TAK-242 (3.0 mg/kg) or *Pg*-LPS (500µg/kg), neither of which influenced blood levels of TNF-alpha when given alone (Fig. 5). Post hoc Sheffé’s tests revealed that the effects of *Ec*-LPS (840 µg/kg) differed significantly from those of vehicle (*P* < 0.01), 500 µg/kg *Pg*-LPS (*P* < 0.01) and 840 µg/kg *Pg*-LPS (*P* < 0.01).

**Fig 5.**
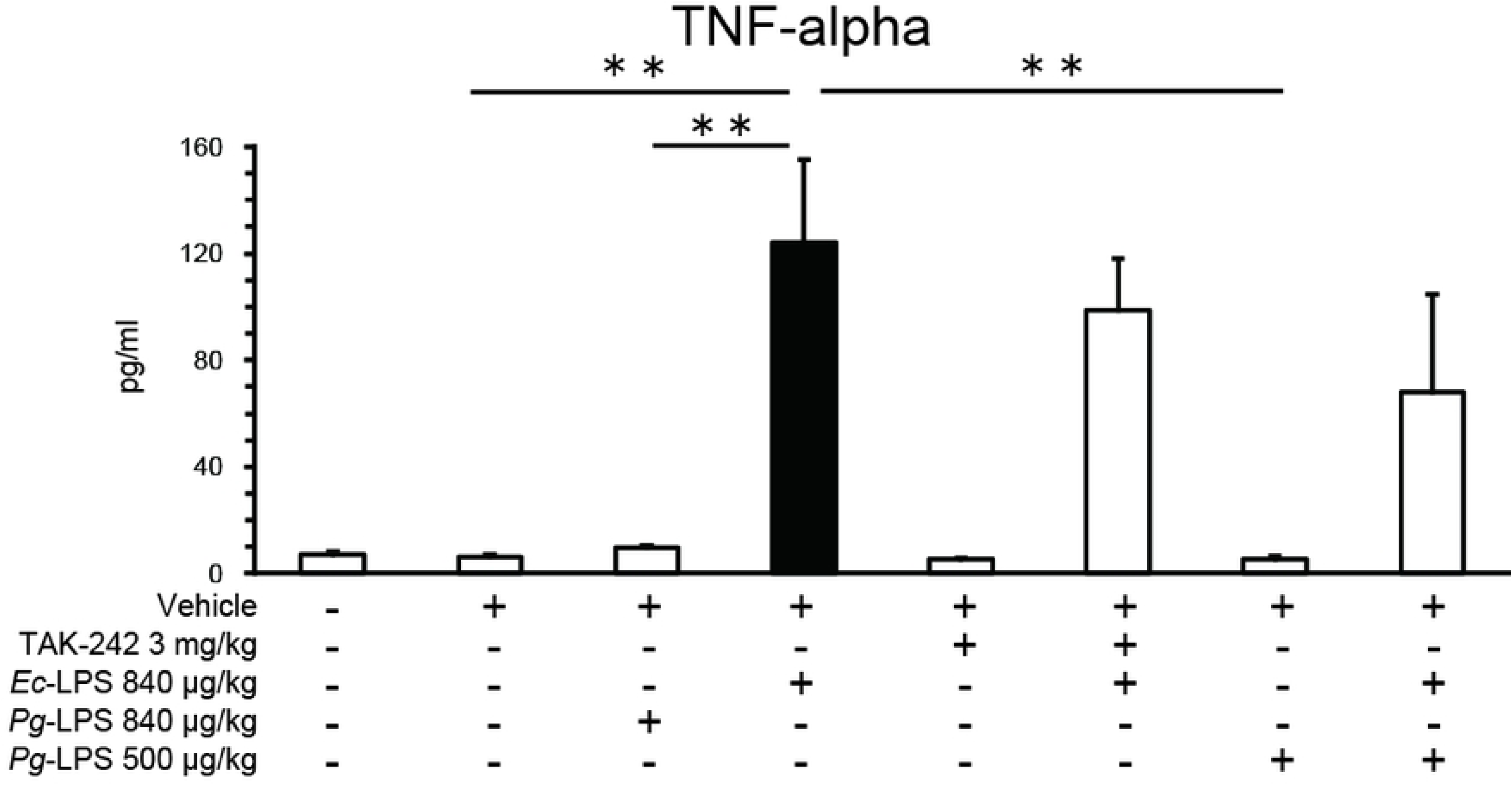
Effects of intra-peritoneal injection of *Pg*- or *Ec*-LPS with TAK-242 and of co-administration of *Pg*- and *Ec*-LPS on TNF-alpha levels (pg/ml) in blood samples of mice. Vertical bars indicate S.E.M., n = 4-6 per group. ** *P* < 0.01 *vs Ec*-LPS + vehicle.

### *Ec*-LPS, but not *Pg*-LPS, increases blood IL-10 levels that are antagonized by TAK-242 and by *Pg*-LPS

Baseline IL-10 levels in blood samples were 91.2 ± 58.7 pg/ml (n = 6) and these levels were not altered by administration of vehicle (Fig. 6). *Ec*-LPS (840 µg/kg), but not *Pg*-LPS (840 ug/kg), increased blood levels of IL-10 (Fig. 6: one-way ANOVA, *F* (7, 34) = 7.22, *P* < 0.001). This *Ec*-LPS (840 µg/kg)-induced increase in blood levels of IL-10 levels was antagonized both by TAK-242 (3.0 mg/kg) and by *Pg*-LPS (500µg/kg), neither of which influenced blood levels of IL-10 when given alone (Fig. 6). Post hoc Sheffé’s tests revealed that the effects of *Ec*-LPS (840 µg/kg) differed significantly from those of vehicle (*P* < 0.01), 500 µg/kg *Pg*-LPS (*P* < 0.01), 840 µg/kg *Pg*-LPS (*P* < 0.01), 3.0 mg/kg TAK-242 + 840 µg/kg *Ec*-LPS (*P* < 0.05), and 500 µg/kg *Pg*-LPS + 840 µg/kg *Ec*-LPS (*P* < 0.05).

**Fig 6.**
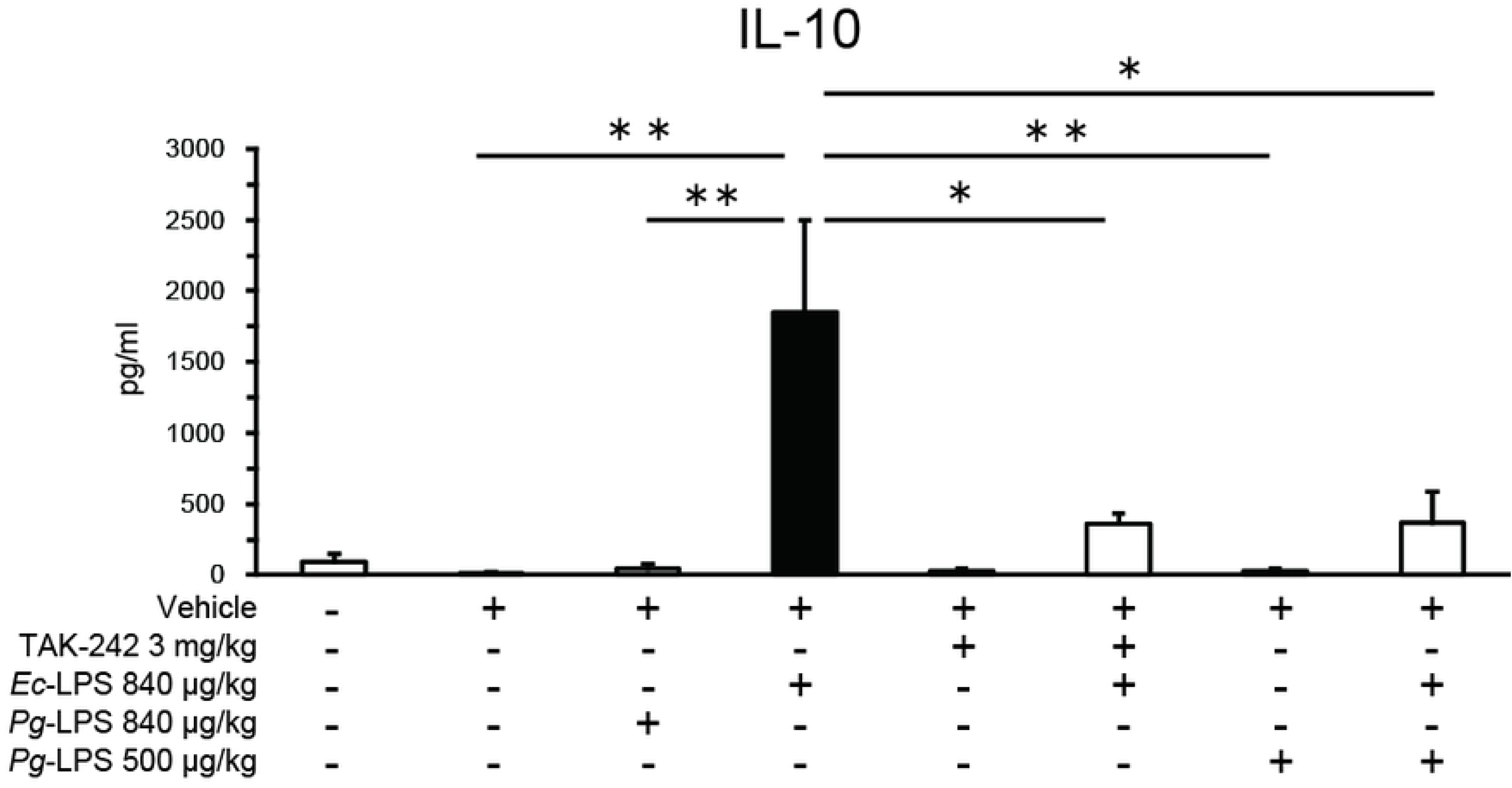
Effects of intra-peritoneal injection of *Pg*- or *Ec*-LPS with TAK-242 and of co-administration of *Pg*- and *Ec*-LPS on IL-10 levels (pg/ml) in blood samples of mice. Vertical bars indicate S.E.M., n = 4-6 per group. \**P* < 0.05, ** *P* < 0.01 *vs Ec*-LPS + vehicle.

## Discussion

The present study indicates that systemic administration of *Ec*-LPS inhibits novel environment-induced increases in locomotor activity by activating TLR4, as this effect was inhibited by the selective TLR4 antagonist TAK-242. To further clarify this *Ec*-LPS-induced inhibition of locomotor activity, more detailed analysis indicated this inhibition to be evident similarly in both the peripheral and central zones of the open field. As anxiety- and/or depression-related processes are reflected in differential effects between these zones [23], such processes are not implicated in the present findings. Rather, they may reflect evidence that LPS-induced reductions in locomotor activity, including spatial exploration, relate to prevention of the spread of inflammation and promotion of healing [16].

In contrast to *Ec*-LPS, systemic administration of *Pg*-LPS did not influence novel environment-induced increases in locomotor activity. Previous findings from our intra-gingival injection study in urethane-anesthetized rats showed that *Pg*-LPS, but not *Ec*-LPS, produced a transient increase in extracellular TNF-alpha levels at the injection site [15]. The present findings provide further evidence that *Pg*-LPS can elicit effects that differ from those of *Ec*-LPS not only in *in vitro* but also *in vivo*. Several *in vitro* studies have indicated that TLR4 can be weakly stimulated [7–10] or inhibited [11–14] by *Pg*-LPS. Based on these findings, it is plausible that the present TLR4-mediated decrease in locomotion induced by *Ec*-LPS might be counteracted by systemic administration of *Pg*-LPS. However, it should be noted that while co-administration of *Pg*-LPS with *Ec*-LPS tended to inhibit *Ec*-LPS-induced changes in locomotion, this failed to attain statistical significance.

It is conceivable that systemic administration of LPS may potentially promote the differentiation of T cells and B cells. This is based on the belief that T cells interact with antigen-presenting cells, including dendritic cells and macrophages that recognize LPS, resulting in their activation and differentiation into specific subtypes, such as helper T cells. Additionally, LPS activates B cells, leading to their differentiation into B cells involved in antibody production, thereby enhancing their ability to produce antibodies. Moreover, systemic administration of LPS is known to induce systemic inflammation and increase spleen weight in experimental animals [18]. However, contrary to these assumptions, the LPS treatments in this study did not increase the number of T or B cells derived from mouse spleen, nor did they increase spleen weight. These results indicate that systemic administration of not only *Pg*-LPS but also *Ec*-LPS could not alter the compositions of lymphocytes, at least within the experimental period of the present studies. Furthermore, the lack of significant changes in spleen weight suggests that these LPS treatments are unlikely to induce material systemic inflammation within this experimental period.

In agreement with an earlier report suggesting that an increase in cytokine synthesis and release mediates the sickness behavior induced by systemic administration of LPS, including decreased novelty-induced locomotion in experimental animals [16], we observed marked increases in blood levels of IL-6, TNF-alpha, and IL-10 in mice treated with *Ec*-LPS. Interestingly, the biological processes involved in the increases in levels of TNF-alpha were not the same as those for IL-6 and IL-10 under the present experimental conditions. Notably, the TLR4 antagonist TAK-242 inhibited *Ec*-LPS-induced increases in blood IL-6 and IL-10 levels but had no effect on TNF-alpha levels. This suggests that IL-6 and IL-10 are likely increased by TAK-242-sensitive TLR4 activation.

Underlying reasons for the failure of TAK-242 to affect *Ec*-LPS-induced increases in TNF-alpha levels are not yet known, but several possibilities may be considered. Firstly, the contribution of intra-cellular TLR4, which may be less accessible to compounds such as TAK-242, could explain its inactivity on TNF-alpha levels [24]. Secondly, differences in TAK-242 concentrations at TLR4 may result in differential blockade, influencing IL-6 and IL-10 levels differently from those of TNF-alpha. For example, under the present *in vivo* conditions TAK-242 might attain a higher concentration at TLR4 sites involved in IL-6 and IL-10 synthesis and release, while attaining a lower concentration at TLR4 sites involved in TNF-alpha synthesis and release. Further investigation would be necessary to clarify these possibilities and gain deeper insights into the mechanisms underlying the differential effects of TAK-242 on cytokine levels.

In contrast to *Ec*-LPS, systemic administration of *Pg*-LPS did not influence novelty-induced locomotion in mice and did not alter basal levels of IL-6, TNF-alpha or IL-10 in blood. These results align with previous *in vitro* findings using gingival fibroblasts, which showed that increases in IL-6 expression induced by *Pg*-LPS were smaller than those induced by *Ec*-LPS [1]. The present results provide *in vivo* biochemical evidence for clear differences in the influences of *Pg*-LPS and *Ec*-LPS on blood IL-6, TNF-alpha and IL-10 levels when administered systemically. Since *in vitro* studies have shown that *Pg*-LPS weakly stimulates [7–10] or inhibits [11–14] TLR4, we investigated the effects of co-administration of *Pg*-LPS on *Ec*-LPS-induced increases in blood IL-6, TNF-alpha and IL-10 levels that appear mediated by TLR4 activation. Distinct from *Ec*-LPS-induced increases in blood TNF-alpha levels, which were not inhibited by TAK-242, *Ec*-LPS-induced increases in blood IL-10, but not IL-6, were suppressed by co-administration of *Pg*-LPS. The reduction in novelty-induced locomotor activity of mice induced by *Ec*-LPS is likely mediated by increases in blood IL-6, as opposed to TNF-alpha and IL-10. This conclusion is supported by the observation that TAK242, but not *Pg*-LPS, counteracted the effect of *Ec*-LPS on mouse locomotion, which corresponded to changes in blood IL-6 levels.

The mechanisms underlying inhibition of *Ec*-LPS-induced increases in blood IL-10 levels by *Pg*-LPS remain unclear. Nevertheless, previous *in vitro* data have suggested that *Pg*-LPS may inhibit the stimulation of TLR4 by *Ec*-LPS [25, 26]. To gain further insights, at least two key aspects require investigation: (1) whether systemically administered *Pg*-LPS acts as a weak agonist or antagonist at TLR4, and (2) the differential effect of *Pg*-LPS treatment on IL-6 and IL-10 levels in response to *Ec*-LPS, considering the observed actions of TAK-242 to counteract these effects on both IL-6 and IL-10. Despite these uncertainties, the present study provides compelling biochemical evidence supporting the efficacy of systemically administered *Pg*-LPS in preventing TLR4-mediated increases in blood IL-10 levels induced by systemic administration of *Ec*-LPS. IL-10 is believed to hinder the progression of periodontitis by regulating mRNA transcription of inflammatory mediators [14] and controlling the production of IFN-gamma and IL-17 by T-cells [27, 28]. Consequently, inhibition of TLR4-mediated increases in IL-10 levels appears to contribute to the development of periodontal diseases triggered by *Pg*-LPS. One of the challenges that should be addressed in future research is any association between inhibition of TLR4, a pattern recognition site involved in the innate immune system, by systemic administration of *Pg*-LPS and induction mechanisms in periodontal diseases and related pathologies other than in periodontal tissue. This is because the effects of *Pg* and *Pg*-LPS are presumed to extend beyond periodontal tissues, potentially influencing various other cell types. For example, *Pg* and *Pg*-LPS have been linked to alterations in neurons within the brain, raising concerns about their potential contribution to the development of Alzheimer’s disease [29]. Similarly, the impact on cardiovascular cells is noteworthy, with possible implications for atherosclerosis [30].

## Conclusion

In summary, this study provides compelling *in vivo* evidence that *Ec*-LPS and *Pg*-LPS exert differential effects on both locomotor activity and cytokine responses in blood. The systemic administration of *Ec*-LPS inhibits novel environment-induced locomotor activity in mice through TLR4 activation. Conversely, *Pg*-LPS failed to affect such locomotion. *Ec*-LPS treatment leads to elevated levels of IL-6, TNF-alpha and IL-10 in blood, likely mediated by TLR4 activation (Il-6 and IL-10) or other pathways distinct from activation of TAK-242-sensitive TLR4 (TNF-alpha). Furthermore, *Pg*-LPS inhibits TLR4-mediated increases in IL-10 levels, while increased levels of IL-6 remained unaffected. The present study provides *in vivo* evidence that *Pg*-LPS can inhibit TLR4-mediated increases in IL-10, which is believed to prevent the development of periodontal disease.

## Abbreviations

LPS: lipopolysaccharide;
*Pg*: *Porphyromonas gingivalis*:
*Ec*: *Escherichia coli*;
TLR: Toll-like receptor;
ANOVA: analysis of variance

## Acknowledgements

The authors would like to express their appreciation for the late Prof. Dr. Mitsuhiro Ohshima, who encouraged us to conduct the present investigations before his passing in 2020.

## Author contributions

Conceptualization of study: H.S., J.L.W., T.S. Formal Analysis: K.S., Y.A., T.H.T., T.S. Funding Acquisition: K.S., Y.A., Y.K., J.L.W., T.S. Investigation: K.S., Y.A., A.W., T.K., T.H.T., T.S. Project Administration: H.O., Y.K., H.S., T.S. Supervision: Y.A., T.H.T., H.S., J.L.W., T.S. Visualization: K.S., Y.A., T.H.T., T.S. Writing – Original Draft Preparation: K.S., T.H.T., Y.K., H.S., J.L.W., T.S. Writing – Review & Editing: K.S., T.H.T., Y.K., H.S., J.L.W., T.S.

## Data availability statement

All data can be obtained upon reasonable request to the corresponding author.

